# The effect of minocycline on the gut microbiome in individuals with alcohol use disorder and healthy controls

**DOI:** 10.64898/2026.10.09.758026

**Authors:** Carolyn M. Doty, Daniel J. Brady, Jennifer J. Barb, Summer L. Thompson, Natalia Rincon, Brittney Browning, Deanna L. Kelly, Melanie E. Bennett, Greg Elmer, Lorenzo Leggio, Brian J. Brandler, Daniel J.O. Roche

## Abstract

Alcohol Use Disorder (AUD) is associated with altered gut microbial composition. Yet, many microbiome studies have included AUD populations with co-occurring conditions (e.g., alcohol associated liver disease; ALD), limiting insight into unique contributions of AUD to gut dysbiosis. Further, preclinical studies show that gut microbiome composition can mediate the behavioral effects of minocycline, including alcohol-related and affective behaviors, but no studies have tested these effects in humans with AUD. This study aimed to 1) compare gut microbiome features between healthy controls (HCs) and individuals with AUD, without ALD, and 2) determine the effects of minocycline on the human gut microbiome. We conducted a four-week randomized, double-blind, placebo-controlled trial of minocycline (100-200 mg/day) in adults with AUD (n=31) and HCs (n=26). We collected fecal samples (n=244) at baseline and each weekly visit for 16S rRNA gene sequencing. Alpha diversity (richness and evenness of taxa within a sample) did not differ by diagnostic group, treatment, or time. In contrast, beta diversity (microbiome composition differences between samples) differed between AUD and controls and was shifted by minocycline over time. AUD, vs. controls, was associated with differences in several taxa prevalences, including reduced *Dialister* and *Subdoligranulum* and increased *Collinsella*, *Streptococcus*, and *Phascolarctobacterium*. Minocycline, vs. placebo, rapidly and robustly restructured the microbiome, altering numerous taxa, including *Lachnospira*, *Ruminococcaceae*, *Clostridium sensu stricto*, and *Christensenellaceae,* and metabolic pathways (e.g., ubiquinol biosynthesis and methylphosphonate degradation). AUD is associated with a distinct gut microbial composition, whereas minocycline produces rapid, sustained, diagnosis-independent gut microbiome restructuring, without altering alpha diversity.

## Introduction

Alcohol Use Disorder (AUD) is a highly prevalent chronic disease, affecting an estimated 28.9 million Americans, and remains one of the leading causes of preventable morbidity and mortality in the United States and globally.^1^ ^2^ Chronic alcohol consumption contributes to more than 200 medical conditions, including liver and cardiovascular diseases, several cancers, neuroinflammation, and immune dysregulation.^3–6^ Despite the well-established burden of AUD, approved pharmacological treatments are very few in number and remain underutilized.^7,8^ Given the heterogeneity of individuals with AUD and the diverse consequences of alcohol use, new approaches to treatment development are urgently needed.

Individuals with AUD exhibit alterations in gut microbial composition.^9–14^ These shifts include depletion of taxa involved in maintaining intestinal barrier integrity^11,15^ and anti-inflammatory functions,^16^ particularly short-chain fatty acid (SCFA)-producing bacteria including *Akkermansia*, *Ruminococcaceae*, *Lachospiraceae*, and *Faecalibacterium prausnitzii.*^17–19^ Furthermore, AUD is associated with enrichment of pathobionts including *Proteobacteria* and *Enterobacteriaceae.*^17,19,20^ In a subset of individuals with AUD, alcohol-induced gut dysbiosis is associated with impaired intestinal barrier integrity, increased intestinal permeability, and systemic inflammation, all of which have been associated with greater alcohol craving and more severe symptoms of depression and anxiety.^9,21–24^ Dysbiosis has been shown to partially normalize with abstinence, indicating that the gut microbiome is dynamically responsive to alcohol exposure and cessation.^14,25^ Furthermore, in individuals with alcohol-associated liver disease (ALD), modulation of the gut microbiome using *Lactobacillus rhamnosus GG* supplementation led to a reduction in liver injury and heavy drinking, highlighting the therapeutic potential of microbiome-targeting therapies^26,27^. Most studies have included cohorts with ALD, individuals in treatment, and/or abstinent individuals, thus limiting insight into the microbiome of individuals with AUD prior to the development of clinically evident liver disease or those not yet engaged in treatment.^10,28^ It therefore remains unclear whether these microbial signatures reflect core features of AUD or, for example, consequences of comorbid clinically evident liver disease.^29^ By comparing individuals with AUD to HCs in a cohort without clinically evident liver disease and spanning the full spectrum of AUD severity, the present study aims to clarify microbiome features that generalize across the disorder and identify potential therapeutic targets.

Minocycline, a broad-spectrum antibiotic with anti-inflammatory and neuroprotective properties, has been shown to alter gut microbial composition and is increasingly studied as a modulator of gut-brain signaling.^30^ As a bacteriostatic tetracycline antibiotic, minocycline suppresses bacterial growth and has the capacity to reshape gut microbial communities.^30,31^ In rodent studies, minocycline consistently alters the gut microbiome, often reducing alpha diversity (the richness and evenness of taxa within a sample) and inducing shifts in community composition, including decreased abundance of Firmicutes and increased Bacteroidetes and Proteobacteria phyla.^32–34^ These changes frequently involve depletion of taxa such as *Lactobacillus* and *Bifidobacterium*, alongside context-dependent increases in families including *Lachnospiraceae* and *Ruminococcaceae.*^32,35,36^ Notably, several rodent models of psychiatric disorders suggest that these minocycline-induced microbiome alterations may contribute to its behavioral effects.^37–40^ Specifically, recent rodent studies suggest minocycline’s effects on the gut microbiome may be *causal* in its anti-depressive, -anxiety, and -alcohol effects and pro-cognitive effects.^32,34,41,42^ Minocycline has also been associated with enrichment of SCFA-producing taxa and improvements in intestinal barrier function in rodents, particularly under conditions of stress or neuroinflammation, supporting a potential microbiome-mediated mechanism for the behavioral and anti-inflammatory effects of minocycline.^35,41,43^

In humans, evidence of how minocycline affects the gut microbiome and its modulation of psychiatric symptoms remains limited and methodologically heterogeneous, with most investigations conducted in clinical populations without healthy control groups and few employing placebo-controlled and within-subjects designs. Such studies suggest that minocycline can alter gut microbial composition, including reductions in *Bifidobacterium* and *Lactobacillus* and increases in *Bacteroidetes*, while effects on alpha diversity are variable, ranging from no change to modest reductions.^44,45^ Clinical trials have explored minocycline for a range of neuropsychiatric conditions, including treatment resistant depression,^46–48^ schizophrenia,^49–51^ and medication-assisted opioid treatment.^52^ These studies yield mixed outcomes: minocycline may improve depressive or cognitive symptoms in some subpopulations, (e.g., those with underlying neuroinflammation), but in larger trials or treatment-resistant cohorts, effects are often non-significant.^46–48,53^ Despite growing interest in microbiome-mediated mechanisms, no studies have directly examined minocycline-induced changes in the gut microbiome of individuals with AUD or analyzed within-subjects changes in HCs. Given preclinical evidence that microbiome alterations may contribute to minocycline’s behavioral effects, we sought to characterize its effects on the human gut microbiome to evaluate potential microbiome-mediated mechanisms.

We conducted a randomized, placebo-controlled clinical trial to assess within-subject changes in the gut microbiome following minocycline administration in individuals with AUD and HCs and to determine whether these microbial effects differed by AUD status. In the parent trial, minocycline did not significantly affect alcohol consumption, craving, or behavioral measures. Microbiome composition was assessed at baseline and at four additional weekly timepoints across four weeks of treatment. Expanding on evidence that AUD is associated with gut microbial alterations and that minocycline can reshape microbial communities in preclinical models, the primary aims of the present study were to characterize differences in gut microbial composition between individuals with AUD and HCs, and to determine whether minocycline induces measurable changes in the human gut microbiome. By leveraging a within-subject design in a cohort spanning the full spectrum of AUD severity, excluding individuals with liver disease, this study seeks to clarify which microbiome alterations are a core feature of AUD and whether minocycline exerts differential microbial effects as a function of AUD status.

## Methods

### Study Overview and Participants

This study was approved by the University of Maryland Baltimore Institutional Review Board, monitored by a Data and Safety Monitoring Board, and conducted in accordance with the Declaration of Helsinki. Participants were recruited from the greater Baltimore area as part of a randomized, double-blind, placebo-controlled clinical trial evaluating the potential of minocycline, a tetracycline-class antibiotic, as a treatment for AUD (NCT04210713).

Eligible participants were between 25 and 65 years of age. Inclusion criteria for the AUD group required a current DSM-5 diagnosis of AUD, weekly consumption of ≥14 drinks for men or ≥7 drinks for women, and ≥5 heavy drinking days in the 30 days prior to screening.^54^ Heavy drinking days were defined as consumption ≥5 drinks for men or ≥4 drinks for women in a single drinking event. Control participants were required to have no current or lifetime DSM-5 psychiatric disorders, consumed <14 or 7 drinks per week for men and women, respectively, in the 30 days before screening, and engaged in ≤ 2 heavy drinking days in the 6 months prior to screening.

Exclusion criteria for all participants included: 1) current or recent (within 30 days) participation in AUD treatment, 2) currently seeking immediate treatment for AUD, 3) current diagnosis of a non-alcohol substance use disorder, 4) current use of psychotropic medications, 5) lifetime diagnosis of schizophrenia or bipolar and related disorders, 6) suicide attempt within the past three years or serious suicidal ideation within the past year, 7) self-reported daily opioid use, 8) clinically significant alcohol withdrawal symptoms (CIWA-Ar >= 10), 9) known hypersensitivity to tetracyclines, 10) currently pregnant or nursing, 11) unwillingness to use reliable contraception during the study, 12) autoimmune or inflammatory conditions that could interfere with safe participation, 13) antibiotic use within the past 30 days, 14) mild traumatic brain injury within the past six months, 15) lifetime history of moderate traumatic brain injury, 16) regular use of prebiotic or probiotic supplements, 17) reading level below the sixth grade level, 18) a positive urine toxicology for cocaine, opioids, amphetamines, methamphetamines, phencyclidine, barbiturates, benzodiazepines, methadone, or tricyclic antidepressants, 19) medical conditions interfering with safe participation (unstable cardiac, renal, liver disease, uncontrolled hypertension, or diabetes), 20) AST, ALT, or GGT≥3x the upper limit of normal, current medication contraindicating minocycline, 21) metal implants/devices incompatible with MRI, 22) participation in a medication clinical trial within the previous 2 months, 23) tested positive for COVID-19 within 3-months of participation, and 24) any other circumstance judged by investigators to compromise participant safety.

### Study Design and Procedures

Participants were recruited from the general population of the greater Baltimore area through online advertisements and social media postings. Participants who responded to these advertisements were contacted to participate in a telephone pre-screening interview. Phone screening involved a description of the nature of the study, verbal consent, and questions covering general health, psychiatric history, and alcohol consumption. Participants deemed eligible were invited to the clinic for an in-person screening visit to provide informed consent for the trial and to confirm eligibility through structured interviews, a physical examination, and clinical laboratory testing.

Following screening and eligibility confirmation, participants with AUD and HCs were randomized to receive either minocycline or a placebo for a 28-day treatment period. Participants assigned to the minocycline group received 100 mg daily during the first week, followed by 200 mg daily for the remaining three weeks. Fecal samples, clinical laboratory assessments (e.g. blood chemistry, liver function tests, and adverse event monitoring), and standardized clinical and behavioral measures were collected at baseline and at four subsequent weekly visits. Dietary intake was assessed using the Rapid Eating Assessment for Participants-Shortened Version (REAP-S) and the Mediterranean Eating Pattern for Americans (MEPA) questionnaires.^55,56^

### Alcohol-Related Measures and Use

Alcohol consumption was assessed using the Timeline Follow-back (TLFB), a validated, calendar-based structured interview designed to capture daily alcohol use patterns.^57^ Participants provided retrospective reports of all alcoholic drinks consumed each day since the last visit, which were used to calculate weekly consumption, frequency of heavy drinking days, and cumulative alcohol intake. We quantified changes in drinking behavior across the study in the AUD group using the TLFB; drinking values for each timepoint in this analysis were recorded as the total drinks consumed in the seven days preceding the study visit. Participants were assessed for psychiatric disorders, including AUD, using the Structured Clinical Interview for DSM-5 (SCID).^58^ Alcohol craving was assessed at each time point using the Alcohol Urge Questionnaire (AUQ): an 8-item self-report instrument that measures an individual’s present craving for alcohol, including desire to drink and expected pleasure from alcohol consumption.^59^

### Fecal Sample Collection and 16S rRNA Sequencing

At each study visit, participants received an at-home fecal sample collection kit to use before their next visit and were instructed to collect the sample as close as possible to that visit. Participants collected their samples in tubes containing DNA/RNA shield and stored them in their home freezer before bringing them to their study visit. Upon receipt, samples were stored at -80°C until processing, and participants received a collection kit for their next visit.

Fecal sample preparation, DNA extraction, and sequencing were performed at the University of Maryland Institute for Genome Sciences (IGS) Genomics Core in two separate batches. DNA was extracted from a ∼100mg sample subaliquot using the MagAttract Power Microbiome DNA/RNA Kit (Qiagen) according to the manufacturer’s protocol and automated on a Hamilton STAR robotic platform. Positive and no-template extraction controls were included in each set of extractions, with preserved bacterial cells serving ass positive extraction controls cells (Zymo). Water was used as the no-template extraction control. The final DNA product was eluted in 100ul molecular grade water and assessed for yield using SYBR green fluorometry prior to being stored at -80°C.

After DNA isolation, a 2-step PCR targeting the V3-V4 region of the 16S rRNA gene was used for DNA library preparation as previously described1 using the Nextera XT DNA Library Kit and unique dual indexing strategy (Illumina). Template DNA was diluted 1:10 prior to the first 16S gene specific PCR. Next-generation sequencing was performed on the Illumina NextSeq 1000 platform using XLEAP 600 cycle, 300bp paired-end read configuration. We used DADA2 to generate amplicon sequence variants (ASVs), and SILVA v132 16S rRNA gene database to train the RDP Naïve Bayesian classifier for taxonomic classification of ASVs.^60–62^

### Statistical Analysis

Analyses were conducted in R (version 4.5.1) using RStudio (version 2025.05.1). To assess changes in drinking behaviors, a linear mixed-effects model was used to test the main effect of week and the treatment-by-week interaction on total drinks consumed and alcohol craving. Microbiome data were processed using the phyloseq package (version 1.48.0).^63^ Only samples with greater than 10,000 reads were included in the analysis. Alpha diversity, which quantifies within-sample taxonomic diversity, was assessed using the number of observed taxa, Shannon Index, and Simpson Index. These indices were calculated using the estimate_richness() function. Longitudinal changes were evaluated relative to baseline (week 0). For each metric, linear mixed-effects models were fitted to account for repeated measures over time using the lme4 R package.^64^ Fixed effects included week, treatment, and diagnostic group, along with their interactions, while sequencing batch, age, and sex were included as covariates. Type III analyses of variance (ANOVAs) were conducted on the fitted models using Satterthwaite’s method to estimate degrees of freedom. Where applicable, post hoc comparisons of estimated marginal means were performed using the emmeans package to examine pairwise differences between time points and groups, with adjustments for multiple comparisons using the Tukey correction.^65^

Beta diversity, reflecting between-sample differences in microbial community composition, was quantified using Aitchison distance, calculated as the Euclidean distance of centered log-ratio (CLR)-transformed OTU counts to account for compositionality.^66^ Permutational multivariate analysis of variance (PERMANOVA) was conducted using the adonis2 function on the Aitchison distance matrix to test for main effects of group, treatment, week and their interactions with sex, age, and sequencing depth as covariates. Repeated measures were accounted for by stratifying permutations by participant ID. To further characterize longitudinal change, distance from baseline was computed for each sample as the Aitchison distance between each follow-up time point and that participant’s baseline sample. These distances were analyzed using linear mixed-effects models including fixed effects of week, treatment, diagnostic group, and their interactions, with a random intercept for participant ID to account for within-subject correlation. Type III ANOVA with Satterwaite’s method was again applied to obtain p-values for fixed effects for the distance from baseline analyses. Post hoc comparisons of estimated marginal means were conducted to examine differences between treatment groups at each time point, with Tukey correction applied to adjust for multiple testing.

We compared the relative abundance and prevalence of taxonomic features at the family and genus levels. MaAsLin3 (version 0.99.18), was used to fit generalized linear models to assess longitudinal changes in the abundance bacterial taxa across the five study visits, while accounting for treatment condition, diagnostic group, and participant-level variation.^67^ Counts were normalized for abundance analyses using total sum scaling and log transformation. MaAsLin3 was also used to convert counts into binary presence or absence values before generating logistic regression models for prevalence comparisons. For each taxon, models were fit that included week, treatment (minocycline vs. placebo), and diagnostic group (AUD vs. control) as fixed effects, along with all two-way and three-way interactions of these variables. Age and sex were included as covariates. To account for within-subject correlation resulting from repeated sampling, the models included a random intercept for participant ID. This allowed us to estimate how taxa abundances and prevalences changed over time, whether these trajectories differed by treatment assignment or diagnostic group, and whether treatment effects varied across weeks or between groups. Taxa with individual q-values < 0.25 were considered significant.

For taxa identified as significant in the MaAsLin3 models, post-hoc evaluations were conducted to determine which weekly time points differed within the significant interaction terms. The fitted MaAsLin3 model objects were imported into R, and estimated marginal means (EMMs) were calculated using the emmeans package. For interactions including week (e.g. week by treatment), EMMs were generated for week within each level of the interacting factor. Pairwise comparisons of weeks were then conducted within each level to identify the specific time points driving the interaction. P-values from these post-hoc contrasts were adjusted for multiple testing using the default Tukey correction implemented in emmeans.

Functional pathway profiles were generated using PICRUSt2 implemented through the Nephele platform. Prior to running PICRUSt2, count data were formatted into BIOM objects and organized in Python (version 3.12) to ensure compatibility with the Nephele workflow. Predicted MetaCyc pathway and Enzyme Commission (EC) abundances obtained from the PICRUSt2 output were imported into R for downstream analysis. Pathway relative abundances were CLR transformed prior to statistical modeling. Longitudinal associations were assessed using MaAsLin3, applying generalized linear models to evaluate changes in pathway relative abundances across five study visits. Fixed effects, interactions, and covariates were identical to those used in the taxa analysis.

## Results

### Sample Characteristics

In the parent study, n = 60 participants attended n = 268 sessions (Table 1). From these visits, we collected n = 263 fecal samples (97.77% of potential fecal samples collected; see Supplementary Table 1). We excluded one sample because the participant provided one sample before the study paused due to the COVID-19 pandemic and later re-enrolled when research resumed. We excluded one participant (n = 5 samples) due to a positive HIV status, one participant (n = 5 samples) due to their acute viral illness during the study, and eight samples due to low read count. We excluded one participant because they withdrew after their baseline visit and their baseline sample did not meet the read count threshold. Thus, n = 244 samples were analyzed in the present study from n = 57 participants. Pill counts and self-report suggest participants took 95.27% of their prescribed medication over the course of the trial, reflecting consistently high adherence (Supplementary Table 2).

**Table 1.**
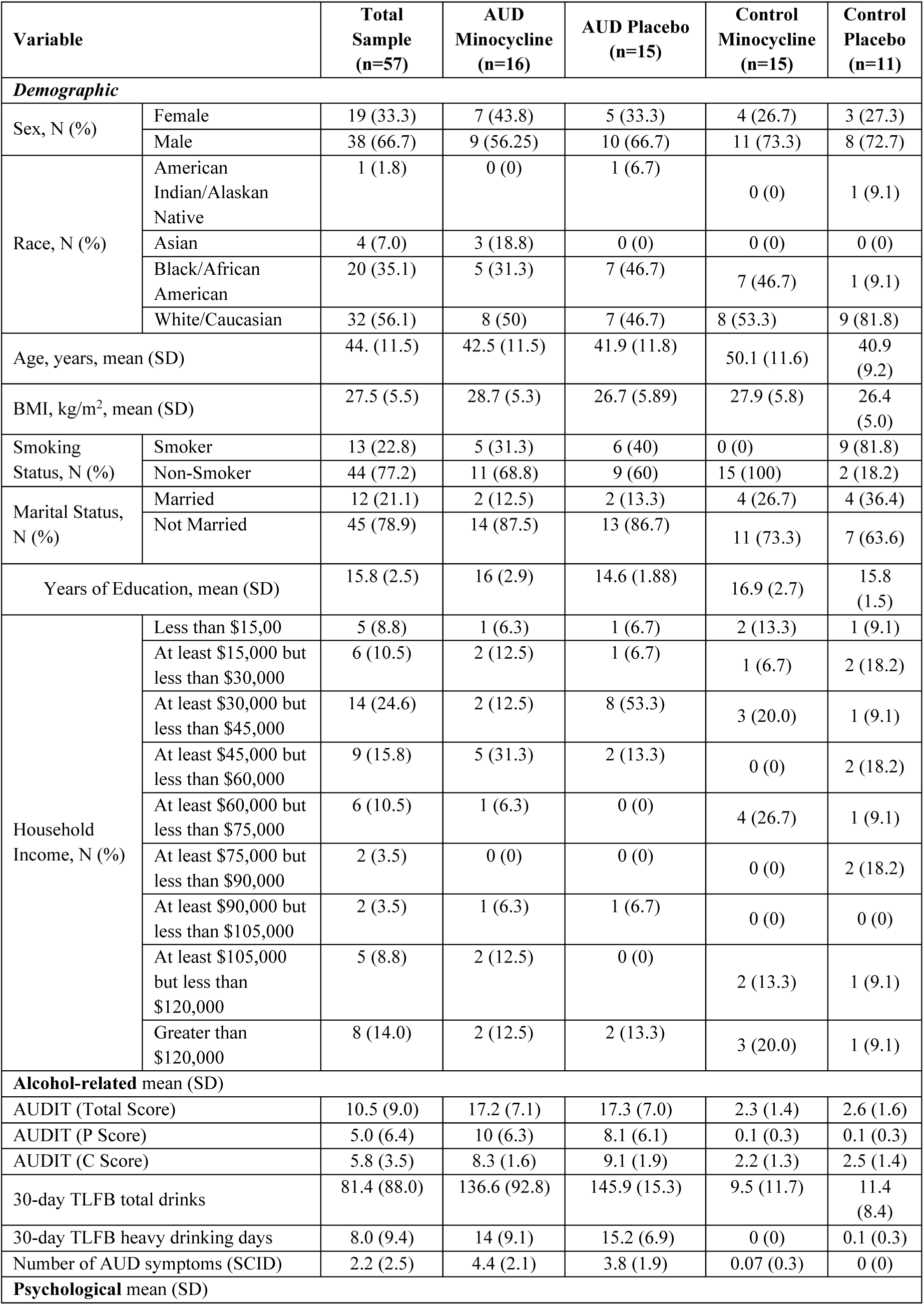
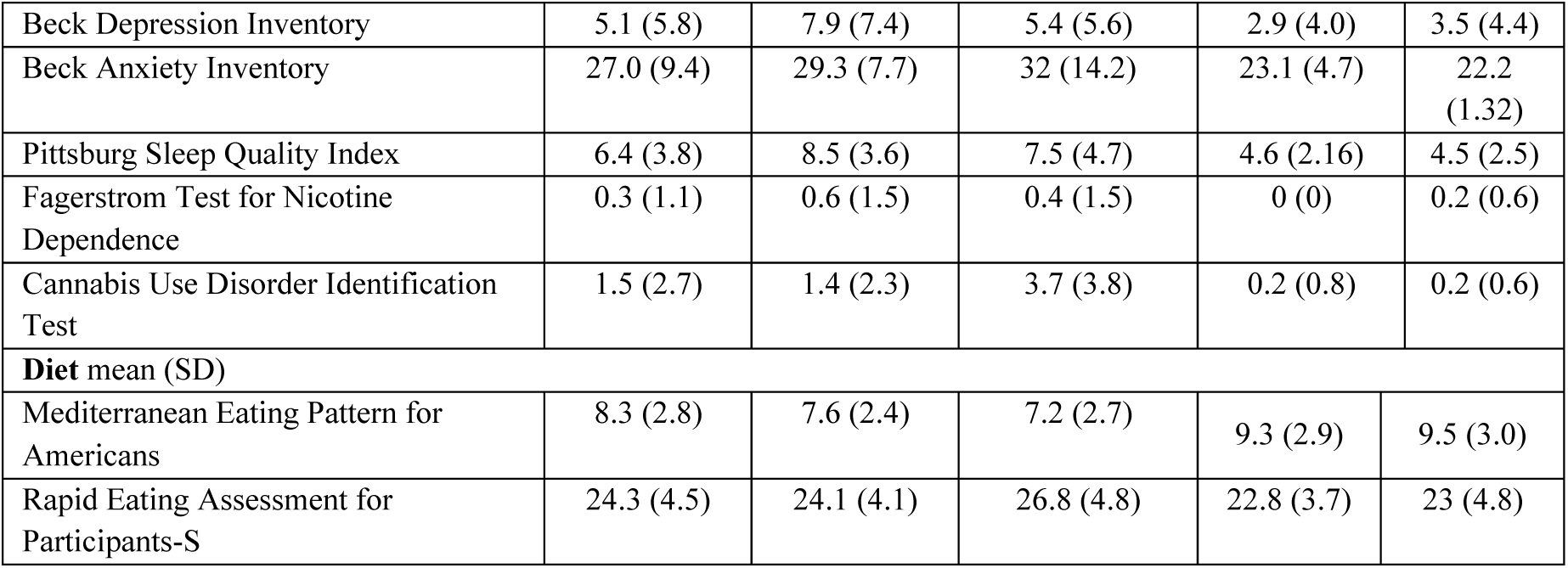
Main Characteristics of Study Population.

| Variable |  | Total Sample (n=57) | AUD Minocycline (n=16) | AUD Placebo (n=15) | Control Minocycline (n=15) | Control Placebo (n=11) |
| --- | --- | --- | --- | --- | --- | --- |
| **Demographic** |  |  |  |  |  |  |
| Sex, N (%) | Female | 19 (33.3) | 7 (43.8) | 5 (33.3) | 4 (26.7) | 3 (27.3) |
|  | Male | 38 (66.7) | 9 (56.25) | 10 (66.7) | 11 (73.3) | 8 (72.7) |
| Race, N (%) | American Indian/Alaskan Native | 1 (1.8) | 0 (0) | 1 (6.7) | 0 (0) | 1 (9.1) |
|  | Asian | 4 (7.0) | 3 (18.8) | 0 (0) | 0 (0) | 0 (0) |
|  | Black/African American | 20 (35.1) | 5 (31.3) | 7 (46.7) | 7 (46.7) | 1 (9.1) |
|  | White/Caucasian | 32 (56.1) | 8 (50) | 7 (46.7) | 8 (53.3) | 9 (81.8) |
| Age, years, mean (SD) |  | 44. (11.5) | 42.5 (11.5) | 41.9 (11.8) | 50.1 (11.6) | 40.9 (9.2) |
| BMI, kg/m<sup>2</sup>, mean (SD) |  | 27.5 (5.5) | 28.7 (5.3) | 26.7 (5.89) | 27.9 (5.8) | 26.4 (5.0) |
| Smoking Status, N (%) | Smoker | 13 (22.8) | 5 (31.3) | 6 (40) | 0 (0) | 9 (81.8) |
|  | Non-Smoker | 44 (77.2) | 11 (68.8) | 9 (60) | 15 (100) | 2 (18.2) |
| Marital Status, N (%) | Married | 12 (21.1) | 2 (12.5) | 2 (13.3) | 4 (26.7) | 4 (36.4) |
|  | Not Married | 45 (78.9) | 14 (87.5) | 13 (86.7) | 11 (73.3) | 7 (63.6) |
| Years of Education, mean (SD) |  | 15.8 (2.5) | 16 (2.9) | 14.6 (1.88) | 16.9 (2.7) | 15.8 (1.5) |
| Household Income, N (%) | Less than \$15,00 | 5 (8.8) | 1 (6.3) | 1 (6.7) | 2 (13.3) | 1 (9.1) |
| | At least \$15,000 but less than \$30,000 | 6 (10.5) | 2 (12.5) | 1 (6.7) | 1 (6.7) | 2 (18.2) |
| | At least \$30,000 but less than \$45,000 | 14 (24.6) | 2 (12.5) | 8 (53.3) | 3 (20.0) | 1 (9.1) |
| | At least \$45,000 but less than \$60,000 | 9 (15.8) | 5 (31.3) | 2 (13.3) | 0 (0) | 2 (18.2) |
| | At least \$60,000 but less than \$75,000 | 6 (10.5) | 1 (6.3) | 0 (0) | 4 (26.7) | 1 (9.1) |
| | At least \$75,000 but less than \$90,000 | 2 (3.5) | 0 (0) | 0 (0) | 0 (0) | 2 (18.2) |
| | At least \$90,000 but less than \$105,000 | 2 (3.5) | 1 (6.3) | 1 (6.7) | 0 (0) | 0 (0) |
| | At least \$105,000 but less than \$120,000 | 5 (8.8) | 2 (12.5) | 0 (0) | 2 (13.3) | 1 (9.1) |
| | Greater than \$120,000 | 8 (14.0) | 2 (12.5) | 2 (13.3) | 3 (20.0) | 1 (9.1) |
| **Alcohol-related mean (SD)** |  |  |  |  |  |  |
| AUDIT (Total Score) |  | 10.5 (9.0) | 17.2 (7.1) | 17.3 (7.0) | 2.3 (1.4) | 2.6 (1.6) |
| AUDIT (P Score) |  | 5.0 (6.4) | 10 (6.3) | 8.1 (6.1) | 0.1 (0.3) | 0.1 (0.3) |
| AUDIT (C Score) |  | 5.8 (3.5) | 8.3 (1.6) | 9.1 (1.9) | 2.2 (1.3) | 2.5 (1.4) |
| 30-day TLFB total drinks |  | 81.4 (88.0) | 136.6 (92.8) | 145.9 (15.3) | 9.5 (11.7) | 11.4 (8.4) |
| 30-day TLFB heavy drinking days |  | 8.0 (9.4) | 14 (9.1) | 15.2 (6.9) | 0 (0) | 0.1 (0.3) |
| Number of AUD symptoms (SCID) |  | 2.2 (2.5) | 4.4 (2.1) | 3.8 (1.9) | 0.07 (0.3) | 0 (0) |
| **Psychological mean (SD)** |  |  |  |  |  |  |

|  |  |  |  |  |  |
| --- | --- | --- | --- | --- | --- |
| Beck Depression Inventory | 5.1 (5.8) | 7.9 (7.4) | 5.4 (5.6) | 2.9 (4.0) | 3.5 (4.4) |
| Beck Anxiety Inventory | 27.0 (9.4) | 29.3 (7.7) | 32 (14.2) | 23.1 (4.7) | 22.2 (1.32) |
| Pittsburg Sleep Quality Index | 6.4 (3.8) | 8.5 (3.6) | 7.5 (4.7) | 4.6 (2.16) | 4.5 (2.5) |
| Fagerstrom Test for Nicotine Dependence | 0.3 (1.1) | 0.6 (1.5) | 0.4 (1.5) | 0 (0) | 0.2 (0.6) |
| Cannabis Use Disorder Identification Test | 1.5 (2.7) | 1.4 (2.3) | 3.7 (3.8) | 0.2 (0.8) | 0.2 (0.6) |
| **Diet mean (SD)** |  |  |  |  |  |
| Mediterranean Eating Pattern for Americans | 8.3 (2.8) | 7.6 (2.4) | 7.2 (2.7) | 9.3 (2.9) | 9.5 (3.0) |
| Rapid Eating Assessment for Participants-S | 24.3 (4.5) | 24.1 (4.1) | 26.8 (4.8) | 22.8 (3.7) | 23 (4.8) |

### Drinking Behavior

Consistent with the parent study, drinking behaviors did not change with treatment in this subsample. No significant effect of week on total drinks consumed (F(4, 90.99) = 0.38, p = 0.824; Supplementary Tables 3 and 4) and no significant week-by-treatment interaction (F(4, 90.99) = 0.78, p = 0.593) in this subsample. Alcohol craving did not significantly change over time (F(4, 87.13) = 2.41, p = 0.056) and showed no week-by-treatment interaction (F(4, 87.13) = 0.49, p = 0.759; Supplementary Tables 5 and 6).

### Alpha Diversity

Observed taxa showed no significant main effects of week, treatment, or diagnostic group, and no significant interactions (See Supplementary Tables 7 and 8). Similarly, Shannon diversity (Supplementary Tables 9 and 10) and Simpson diversity (Supplementary Tables 11 and 12) indices showed no significant main effects or interactions (Figure 1).

**Figure 1.**
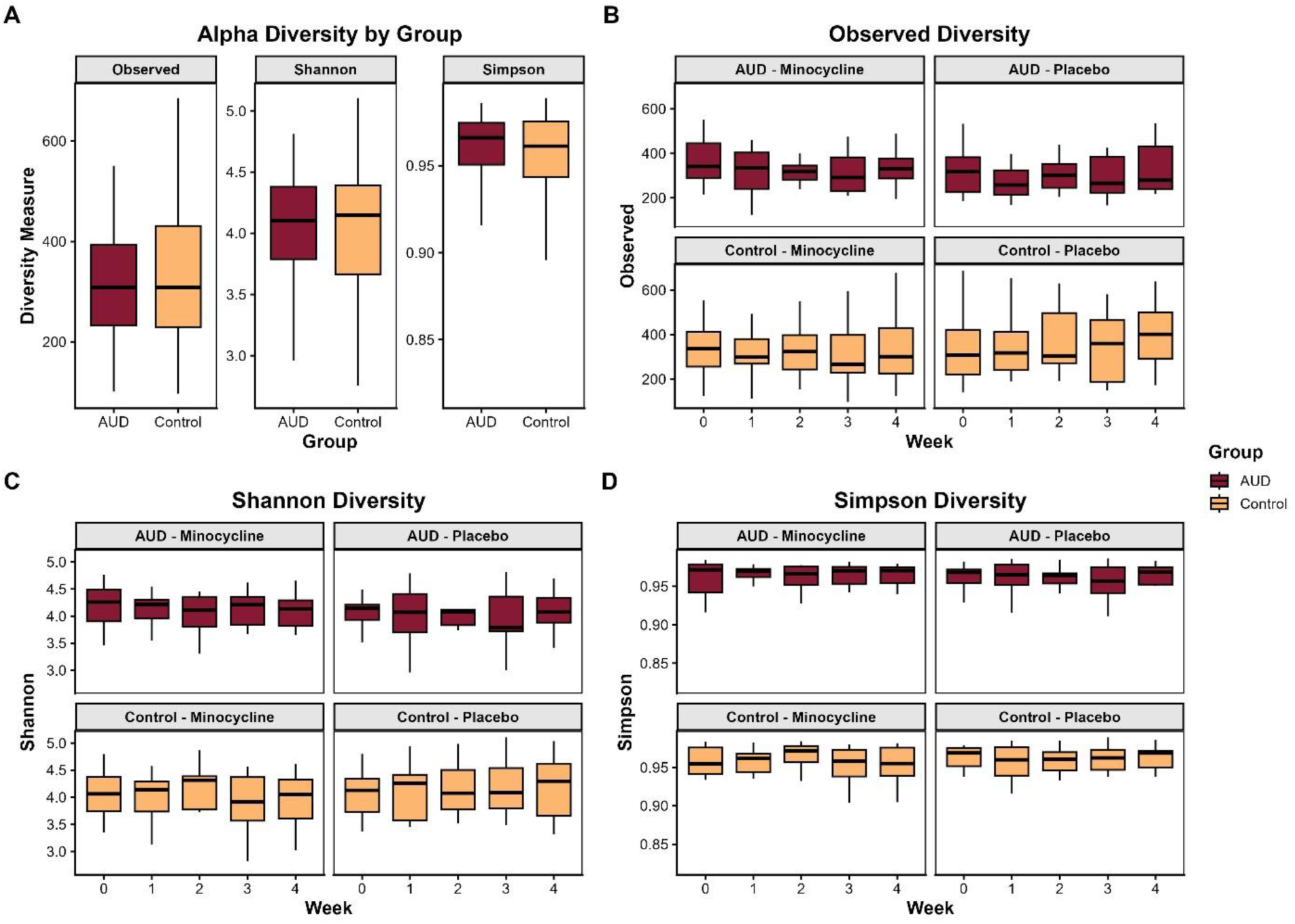
Gut microbiome alpha diversity across clinical and treatment groups. The center lines represent the median, the box represents the interquartile range, and whiskers represent values within 1.5 x the interquartile range. **a.** Alpha diversity comparisons between participants with alcohol use disorder (AUD) and healthy controls using observed richness, Shannon diversity, and Simpson diversity indices. Samples from all timepoints were pooled for comparison. **b.** Longitudinal changes in observed richness across the 5-week study period, stratified by diagnosis and treatment groups. No significant longitudinal changes were observed. **c.** Longitudinal changes in Shannon diversity across diagnosis and treatment groups. No significant longitudinal changes were observed. **d.** Longitudinal changes in Simpson diversity across diagnosis and treatment groups. No significant longitudinal changes were observed.

### Beta Diversity

PERMANOVA revealed main effects of diagnostic group (R^2^ = 0.02, F(1, 221) = 4.37, p<0.001), treatment (R^2^ = 0.02, F(1, 221) = 4.45, p<0.001), and week (R^2^ = 0.01, F(4, 221) = 0.41, p<0.001), and significant interactions for group-by-treatment (R^2^ = 0.01, F(1,221) = 3.50, p<0.001) and treatment-by-week (R^2^ = 0.01, F(4,221) = 0.40, p<0.001). Sex (R^2^ = 0.03, F(1, 221) = 7.15, p<0.001), age (R^2^ = 0.02, F(1, 221) = 3.99, p = 0.032), and sequencing depth (R^2^ = 0.01, F(1,221) = 3.77, p<0.001) also exhibited main effects (See supplementary Table 13)(Figure 2).

**Figure 2:**
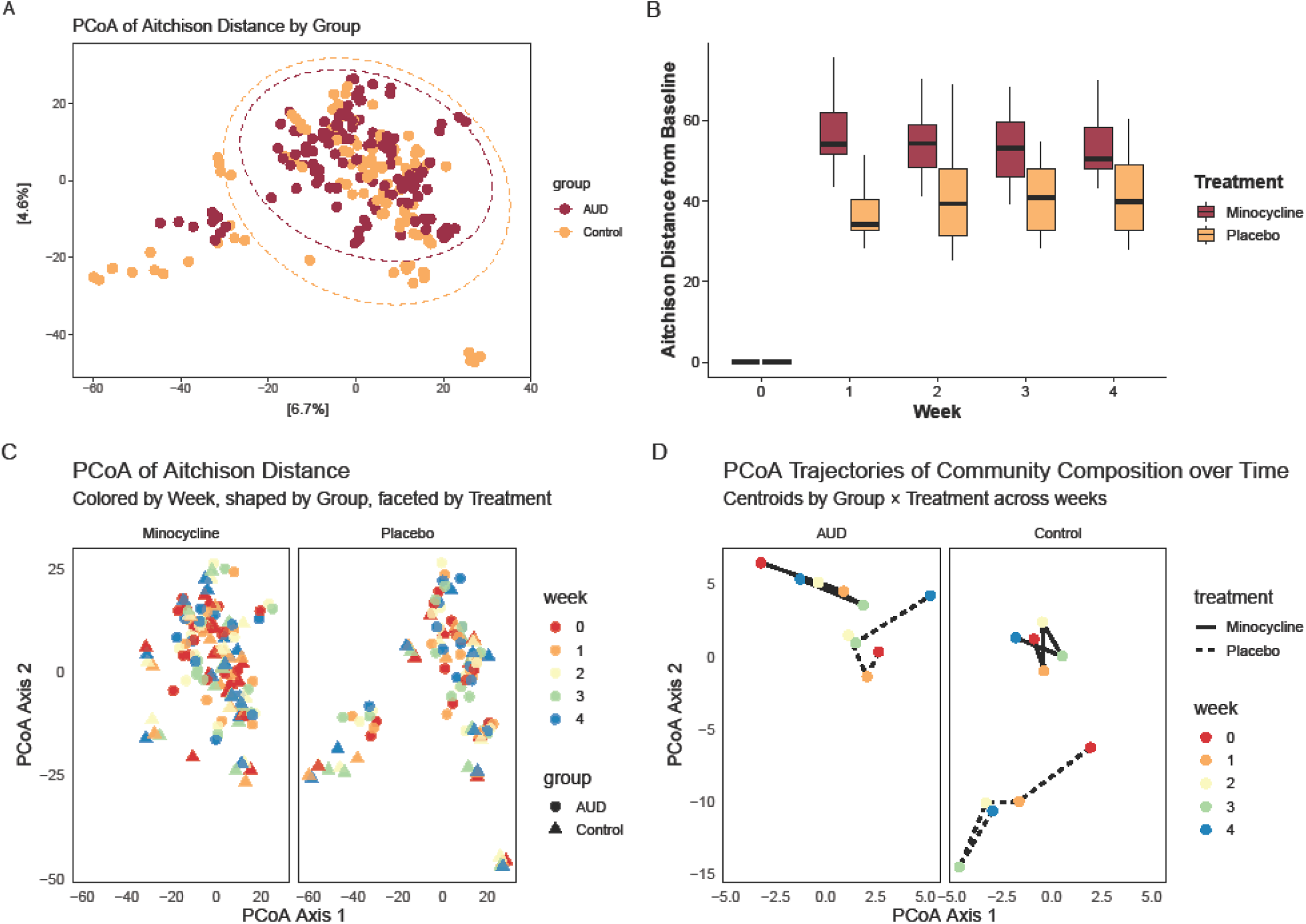
Gut microbiome beta diversity across diagnostic groups, treatment groups, and time. **a.** Principle coordinates analysis (PCoA) of Aitchison distances derived from centered log ratio transformed counts from participants with alcohol use disorder (AUD) and controls. Microbial community composition differed significantly between groups. **b.** Aitchison distance from baseline across the 5-week clinical trial stratified by treatment group. Distances were calculated relative to each participant’s baseline sample (week 0). Samples from participants who were administered minocycline had a greater distance from baseline than those who were administered placebo. The center lines represent the median, the box represents the interquartile range, and whiskers represent values within 1.5 x the interquartile range. **c.** PCoA of Aitchison distances colored by study week and shaped by diagnostic group with panels faceted by treatment group. There were significant effects of diagnostic group, treatment group, and a significant treatment-by-week interaction. **d.** Temporal trajectories of group centroids in ordination space across study weeks for each diagnostic and treatment group combinations. Points represent week-specific centroids and lines indicate trajectories over time.

Distance-from-baseline analyses (assessed by Aitchison distance) showed significant effects of week (F(4, 173) = 688.23, p <0.00001), treatment (F(1,51) = 37.37, p<0.00001), and their interaction (F(4, 173) = 17.91, p<0.00001)(Figure 2; Supplementary Tables 14 and 15). Pairwise comparisons indicated greater distance from baseline in the minocycline group relative to placebo at all timepoints (all p<0.0001)(Supplementary Tables 16 and 17).

### Taxa comparisons

At a false discovery rate threshold of q<0.25, MaAsLin3 revealed multiple taxa which showed significant main effects of diagnostic group, two-way treatment-by-week interactions, and three-way group-by-treatment-by-week interactions for prevalence. Eight taxa exhibited significant treatment-by-week interactions for relative abundance (Supplementary Tables 18 and 19).

#### Diagnostic Group Effects

No taxa showed significant differences in relative abundance between individuals with AUD and controls while controlling for week, treatment and their interactions (Supplementary Tables 18 and 19). By contrast, AUD was associated with significant differences in taxon prevalence, with seven taxa showing increased prevalence and two taxa showing decreased prevalence relative to controls (Figure 3). Specifically, the prevalence of *Dialister* and *Subdoligranulum* was reduced in AUD, whereas *Collinsella*, *Coriobacteriaceae*, *Ruminococcaceae_UCG-*004, *Phascolarctobacterium, Agathobacter, Acidaminococcaceae,* and *Streptococcus* showed increased prevalence (Supplementary Tables 20 and 21).

**Figure 3.**
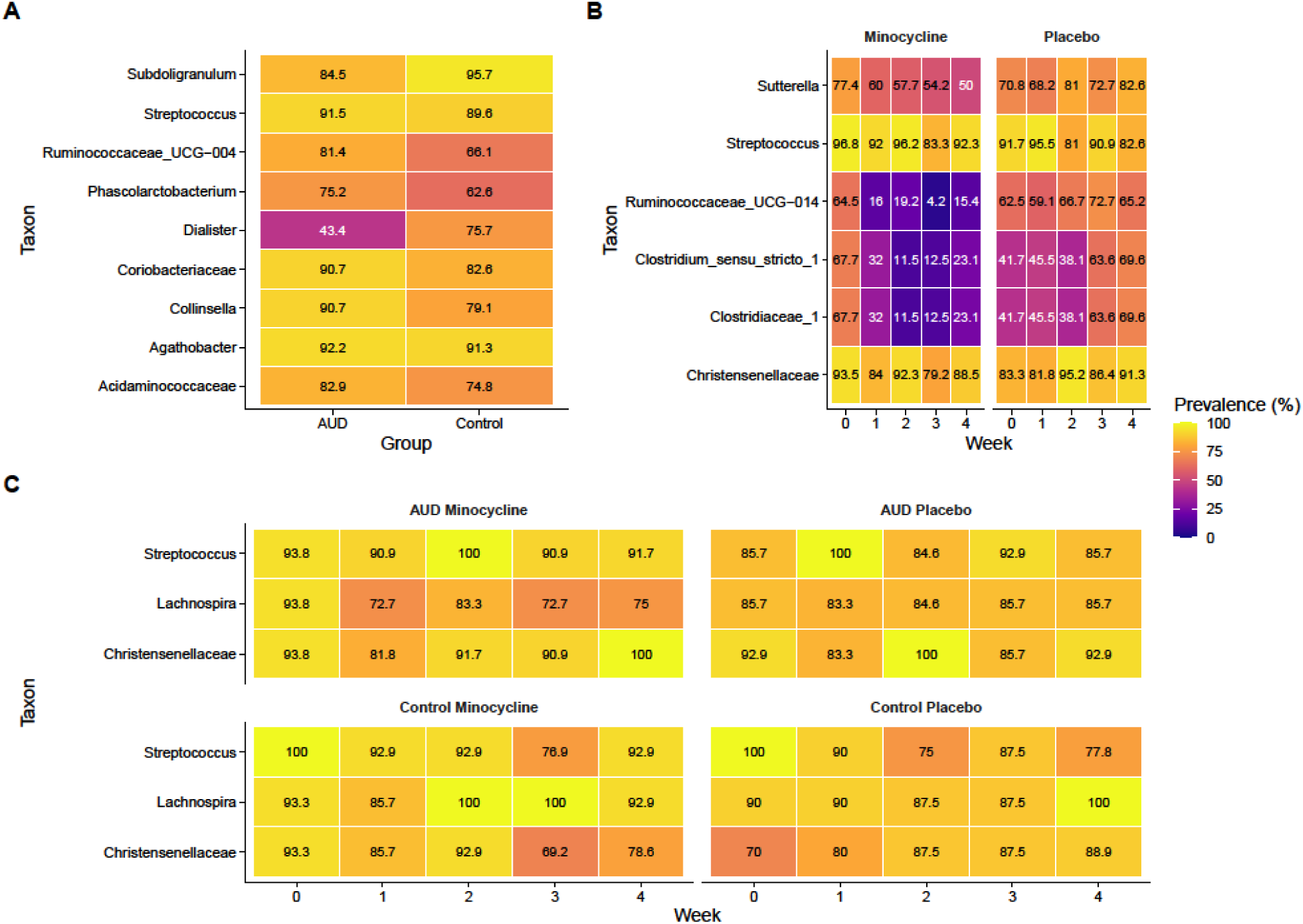
Differential prevalence of gut bacterial taxa across study visits, diagnostic groups, and treatment conditions. **a.** Heatmap showing the prevalence of taxa identified as significantly different between participants with alcohol use disorder (AUD) and controls using MaAsLin3. Tile color indicates prevalence (%) within each diagnostic group, and values within tiles denote the corresponding prevalence percentage. **b.** Heatmap showing the prevalence of taxa identified as significantly associated with treatment conditions across the 5-week clinical trial. Taxa prevalence is shown across study weeks and faceted by treatment condition. **c.** Heatmap showing the prevalence of taxa identified as significantly associated with he interaction between diagnostic group, treatment condition, and study week. Panels are stratified by group-treatment combinations.

#### Treatment Group Effects

Among participants receiving minocycline, several taxa demonstrated consistent decreases in prevalence over time, including *Clostridium sensu stricto 1*, *Ruminococcaceae UCG-14*, and *Clostridiaceae 1* (Figure 3)*. Streptococcus* prevalence decreased over time in the minocycline group, whereas changes in the placebo group were less consistent across timepoints. *Suterella* showed reduced prevalence during weeks 2-4 in the minocycline group. In contrast, *Christensenellaceae* prevalence increased over time within both treatment groups (Figure 3; Supplementary Tables 24 and 25).

Minocycline treatment was associated with decreases in the relative abundance of *Dialister* and *Desulfovibrio*, and increases in *Flavonifractor* and *Parasutterella* at specific timepoints. *Enterobacteriaceae* showed decreased relative abundance in the minocycline group over time. Three additional taxa *Faecalitalea, Streptococcus,* and *Solobacterium* had significant treatment-by-time interactions but did not reveal significant treatment effects at any point in pairwise contrasts (Figure 4; Supplementary Tables 26 and 27).

**Figure 4.**
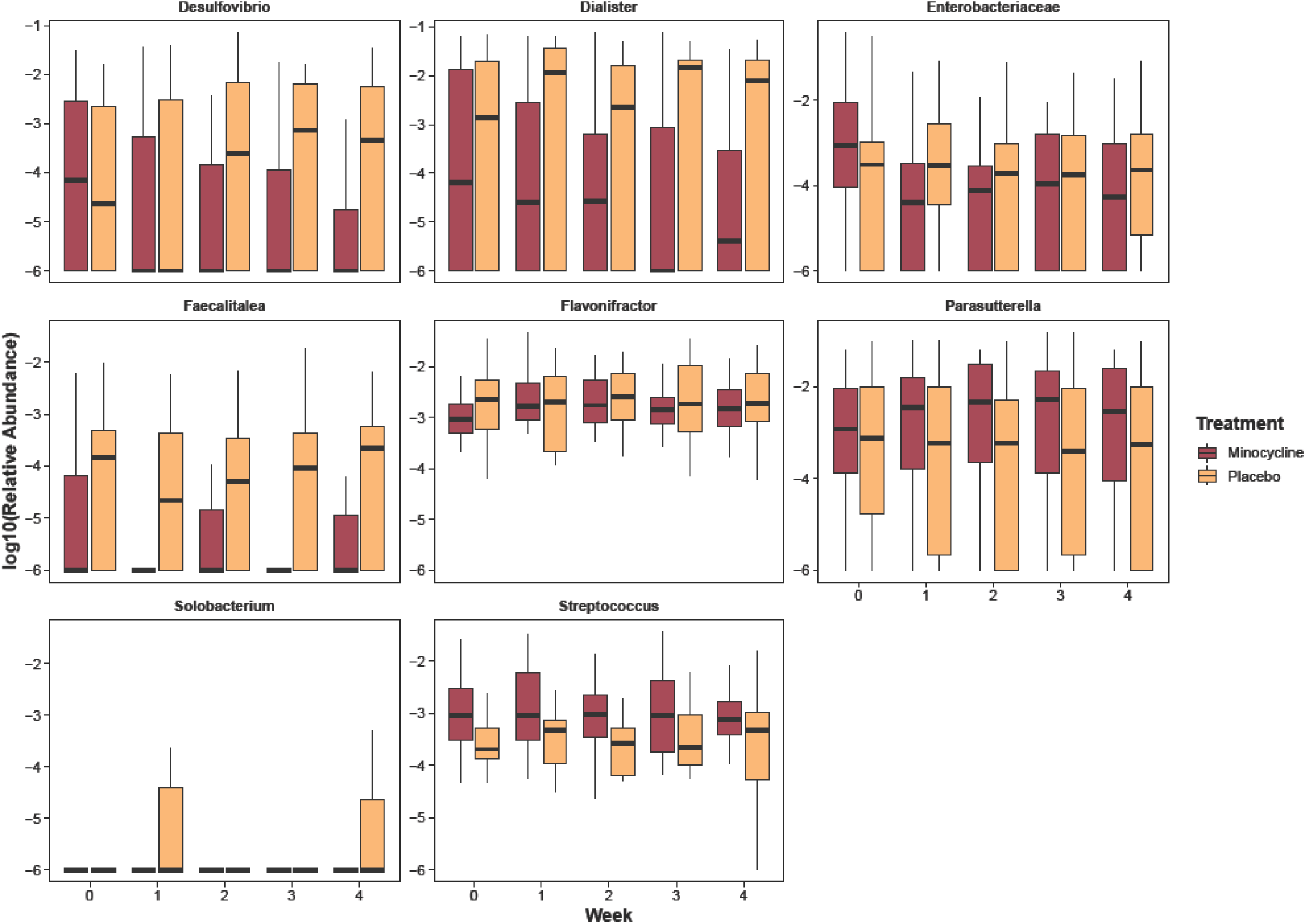
Differential abundances of bacterial taxa associated with treatment condition-by-week interaction. Log10-transformed relative abundances of significantly associated bacterial taxa at the family and genus levels across the 5-week clinical trial. Abundances are shown across study weeks, with week – representing baseline prior to medication administration. Data are shown as boxplots, with boxes representing the interquartile range and horizontal lines indicating medians; whiskers extend to 1.5x the interquartile range. Panels are faceted by taxon. Only data identified as significantly associated with the treatment-by-week interaction are shown

#### Diagnostic Group-by-Treatment-by-Time Interactions

Three taxa prevalences exhibited significant group-by-treatment-by-time interactions. *Lachnospira* prevalence decreased over time in AUD participants receiving minocycline and increased in control participants in both the placebo and minocycline conditions. *Streptococcus* exhibited a significant three-way interaction with prevalence increasing transiently in AUD participants administered minocycline at week 2 and at week 1 in the placebo-AUD group, while decreasing significantly across all weeks in both control groups. Relative to baseline, *Christensenellaceae* prevalence significantly increased at week 4 in the minocycline-treated AUD group and at week 2 in the placebo-AUD group, whereas no significant longitudinal changes were detected in control participants (Figure 3; Supplementary Tables 22 and 23).

### Functional Pathways

A total of 24 MetaCyc pathways showed significant treatment-by-week interactions (q<0.25), including four pathways involved in ubiquinol biosynthesis (Figure 4; Supplementary Table 28). One pathway (methylphosphonate degradation I) showed a significant three-way interaction between diagnostic group, treatment, and week. No pathways were significantly associated with diagnostic group alone (Figure 5).

**Figure 5.**
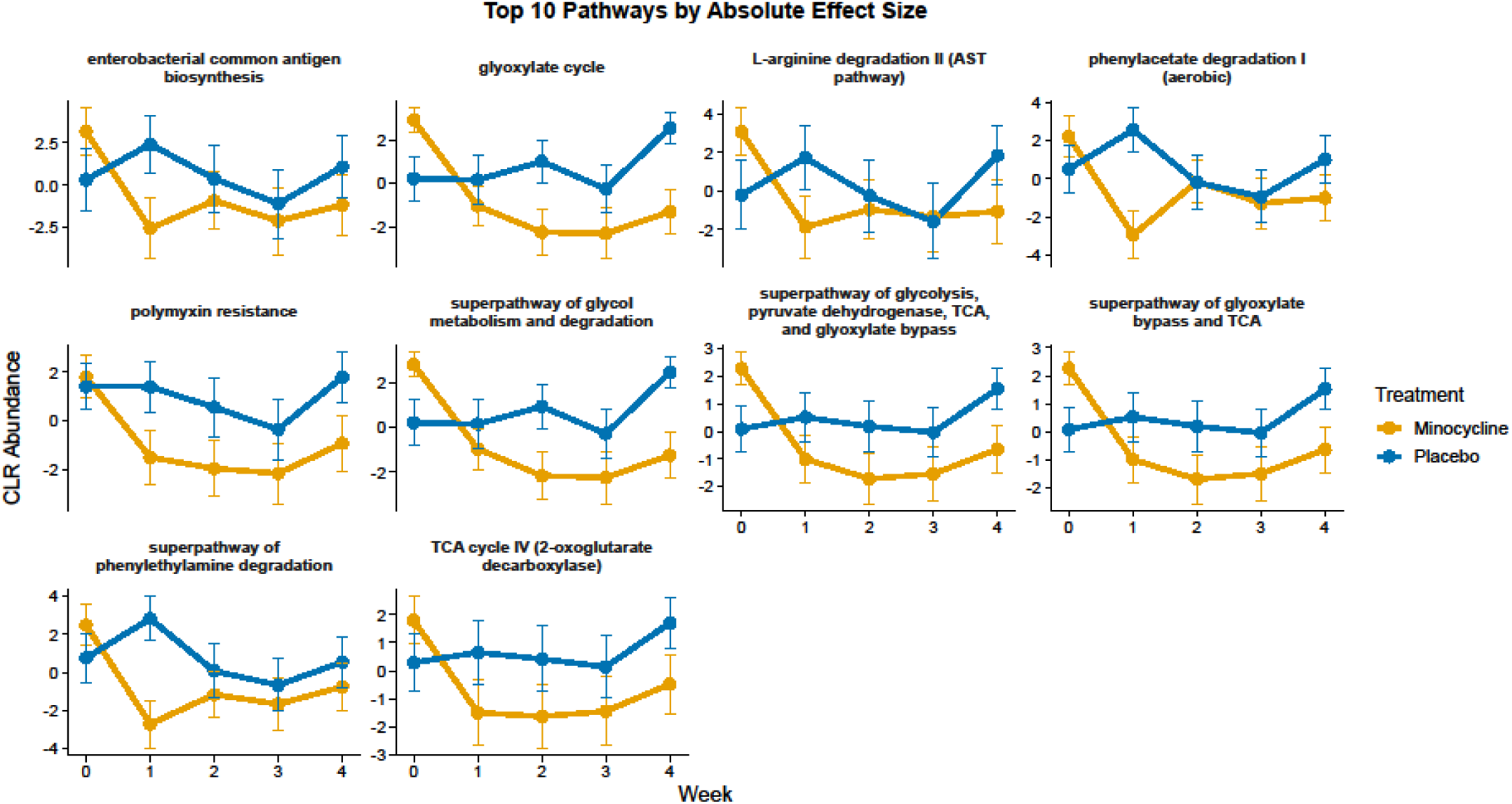
Longitudinal dynamics of predicted MetaCyc pathways associated with treatment effects. Centered log-ratio transformed abundances of the top 10 significantly associated MetaCyc pathways identified with PICRUSt2-predicted functional profiles, ranked by absolute effect size across the 5-week clinical trial. Longitudinal trajectories are shown across study weeks (week 0 denotes baseline prior to treatment initiation). Lines represent group-level mean trajectories, points indicate mean values, and error bars represent ± one standard error of the mean.

Diagnostic group was significantly associated with 10 Enzyme Commission (EC) numbers. A total of 163 EC numbers showed significant treatment-by-week interactions and 26 EC numbers showed significant group-by-treatment-by-week interactions (Supplementary Table 29). Notably, several pathways involved in amino acid metabolism had significant treatment-by-week interactions including lysine decarboxylase, taurine dioxygenase, and 4-aminobutyrate transaminase.

## Discussion

In this secondary analysis of a randomized, placebo-controlled trial, this study was the first to evaluate the effects of minocycline on the gut microbiome of individuals with AUD and HCs. Consistent with the parent trial, minocycline did not significantly alter alcohol consumption or craving in this sub-sample. Alpha diversity measures remained stable across all time points, treatment groups, and diagnostic groups, indicating no changes in taxa richness or evenness over the study period. Conversely, we observed modest differences in overall microbial composition between diagnostic groups, with more pronounced differences at the taxon level where several genera showed altered prevalence in individuals with AUD despite no differences in alpha diversity. Beta diversity analysis revealed robust shifts in microbial composition associated with minocycline treatment independent of diagnostic group. These shifts were particularly apparent when comparing Aitchison distance from baseline, where the minocycline group exhibited greater changes in microbial composition than the placebo group at each time point. These findings indicate that minocycline restructures microbial community composition without altering richness, evenness, or short-term behavioral outcomes.

Differences observed between individuals with AUD and HCs are associated with taxa that are implicated in metabolic regulation, inflammation, and gut-brain signaling. Compared to controls, participants with AUD exhibited reduced prevalence of *Dialister*, a propionate-producing genus, previously linked to psychiatric conditions including major depressive disorder and schizophrenia.^68–70^ Among other SCFA-producers, increased prevalence of *Coriobacteriaceae* and reduced prevalence of *Sudoligranulum* were consistent with prior findings in heavy drinking populations.^71–73^ The observed increase in *Acidaminococcaceae* is notable because a previous study found reduced abundance in heavy-drinking individuals, suggesting a potential microbiome shift during the transition from heavy drinking to clinically significant AUD.^74^ AUD was also associated with increased *Streptococcus*; elevated *Streptococcus* has been associated with worse alcohol-associated liver disease, major depressive disorder, and schizophrenia.^75,76^ The increased prevalence of *Collinsella* is notable as this genus has been associated with pro-inflammatory signaling and elevated cytokine levels, is enriched in psychiatric conditions such as schizophrenia as well as in individuals exhibiting post-traumatic stress symptoms, and has been associated with low fiber diets.^77–80^ Collectively, these findings indicate that AUD is associated with a microbial profile marked by reduced SCFA-producing taxa and enrichment of pro-inflammatory bacteria, consistent with a shift toward a more inflammatory gut environment. These findings are consistent with prior studies demonstrating gut microbial dysbiosis in AUD and further support the gut microbiome as a dysregulated biological system in AUD. Overall, the microbial profile observed in individuals with AUD is characterized by taxa associated with inflammatory, metabolic, hepatic, and psychiatric disorders, suggesting a gut environment that may contribute to AUD and/or ALD pathology and medical conditions.

Our findings indicate that individuals with AUD exhibited a distinct microbial community structure relative to controls. These differences were primarily observed in taxon prevalence rather than relative abundance, suggesting that AUD may be characterized by inconsistent presence of specific less-abundant microbes across individuals rather than coordinated changes in dominant taxa. This distinction is biologically meaningful as prevalence-based differences may reflect disruptions in microbial resilience or host-microbe compatibility after repeated alcohol exposure, rather than expansion or depletion of dominant taxa.^81^ These findings, taken together with previous research demonstrating that supplementation of beneficial bacterial taxa, specifically *Lactobacillus rhamnosus GG*, associated with improvements in liver function and reductions in alcohol function, support the gut microbiome as a promising potential therapeutic target for AUD.^26^

Minocycline induced selective restructuring of the gut microbiome, characterized by consistent reductions in several SCFA-producing taxa alongside enrichment of taxa implicated in metabolic and immune processes. Notably, this included a reduction in *Dialister*, a propionate-producing genus linked to mental health outcomes, that was already reduced in individuals with AUD. Decreases in *Lachnospira*, *Clostridium sensu stricto 1*, and *Ruminocacceae UCG-014* suggest that minocycline may reduce capacity for SCFA production, with potential downstream effects on gut barrier integrity and host metabolism.^82–84^ By contrast, increases in *Christensenellaceae*, a taxon associated with metabolic health and reduced risk of affective disorders, point to parallel shifts that may reflect compensatory or context-dependent microbial responses.^85,86^ Changes in *Streptococcus*, a genus often exhibiting tetracycline resistance, were transient and differed by diagnostic group, underscoring the heterogeneity in antibiotic susceptibility.^87^ Additional alterations, including reductions in *Dialister* and *Desulfovibrio* and increases in *Flavonifractor* and *Parasutterella*, further indicate that minocycline impacts taxa involved in microbial metabolism, bile acid regulation, and immune signaling.^88–90^ Notably, the decrease in *Enterobacteriaceae* abundance in our participants contrasts with a prior report of enrichment following tetracycline exposure using an *in vitro* model, suggesting that host context may shape antibiotic response.^91^ Additionally, *Lachnospira*, *Streptococcus*, and *Christensenellaceae* exhibited distinct transient longitudinal trajectories across treatment and diagnostic groups, suggesting that baseline microbial ecology in AUD may influence gut bacterial responses to minocycline. Additionally, predicted functional pathway analysis using PICRUSt2 revealed treatment-by-week interactions in pathways including ubiquinol biosynthesis, suggesting that minocycline-induced compositional changes may influence antioxidant production and energy metabolism. Together, these findings indicate that minocycline exerts targeted ecological pressure on the gut microbiota, preferentially altering metabolically and immunologically relevant taxa without affecting overall diversity.

This study should be viewed in the context of its strengths and limitations. In the absence of detectable clinical effects of minocycline, we could not link microbiome changes to behavioral outcomes, suggesting that microbiome changes did not translate into short-term behavioral effects. However, it is possible that a 4-week trial was simply too short to detect behavioral effects of minocycline related to changes in alcohol consumption or craving. As our observed microbiome shifts were greatest at week 4, it is plausible that microbiome composition would have continued to change with further treatment and that these microbial alterations may *precede* behavioral changes that occurred outside of the study window. In addition, taxa did not respond linearly nor incrementally across time, as some exhibited rapid changes early in treatment, whereas others shifted more gradually. As diet has a profound impact on the gut microbiome and AUD is associated with malnutrition, this study is limited by the lack of rigorous diet measures.^92^ Future studies should incorporate longer intervention durations, post-treatment follow-up, and more frequent sampling to determine whether microbiome changes persist, normalize, or ultimately translate into delayed behavioral or clinical outcomes. This study, to our knowledge, is the first to examine the within-subject impact of minocycline on the gut microbiome while including a control group, enabling a clearer separation of treatment versus diagnostic effects, which is particularly important as minocycline is a frequently prescribed medication. The longitudinal repeated-measures design further allowed us to characterize within-subject microbiome dynamics across the intervention period.

In summary, AUD was associated with distinct gut microbial alterations that may impact SCFA production, inflammation, and metabolic function, reinforcing the growing body of evidence that the gut microbiome may contribute to the pathophysiology of AUD and alcohol-related medical conditions. Although preclinical studies have suggested that minocycline may provide therapeutic benefits through modulation of the gut microbiome, our findings did not indicate that minocycline restored AUD-associated microbial alterations or selectively normalize the gut microbiome in participants with AUD. Instead, minocycline produced substantial restructuring of the gut microbiome regardless of diagnostic group, raising the possibility that this treatment regimen introduces additional microbial perturbations rather than correcting existing dysbiosis. These findings are clinically relevant because minocycline is widely prescribed, often for prolonged or repeated courses, yet its effects on the gut microbiome remain incompletely understood.^30^ Even in the absence of immediate behavioral effects during the four-week trial, minocycline-induced changes in microbial ecology may have downstream consequences for gut and metabolic health, particularly in populations who already exhibit microbiome dysbiosis, such as individuals with AUD. Future studies should determine whether these microbial alterations persist after treatment attrition, define their functional consequences, and establish whether longer treatment periods reveal delayed effects on alcohol-related or other clinically relevant outcomes.

## Funding

This study was supported by NIAAA K01AA026005 (D.J.O.R).

## Disclosure

LL is supported by the NIH IRP (NIDA and NIAAA). The contributions of the NIH author are considered Works of the US Government. The findings and conclusions presented in this paper are those of the authors and do not necessarily reflect the views of the NIH or the US Department of Health and Human Services.

### COI

DLK has served on an advisory board for Amani Therapeutics, Boehringer Ingelheim and Teva. She served as a consultant for Alkermes. Outside of his NIH work, Dr. Leggio receives an honorarium from UK Medical Council on Alcohol as Editor-in-Chief for Alcohol and Alcoholism, and he also receives royalties from Rutledge as co-editor of a textbook.

## Data availability

Data are currently not publicly available but will be made available upon request.

## References

1. Alcohol-Related Disease Impact | CDC. https://nccd.cdc.gov/DPH_ARDI/default/default.aspx.

2. WHO. Global status report on alcohol and health and treatment of substance use disorders. https://www.who.int/publications/i/item/9789240096745.

3. Alcohol and Cancer Risk Fact Sheet - NCI. https://www.cancer.gov/about-cancer/causes-prevention/risk/alcohol/alcohol-fact-sheet (2021).

4. Sahu, P., Verma, H. K. & Bhaskar, L. Alcohol and alcoholism associated neurological disorders: Current updates in a global perspective and recent recommendations. World J. Exp. Med. 15, 100402 (2025).

5. Biddinger, K. J. et al. Association of Habitual Alcohol Intake With Risk of Cardiovascular Disease. *JAMA Netw*. Open 5, e223849 (2022).

6. Szabo, G. & Saha, B. Alcohol’s Effect on Host Defense. Alcohol Res. Curr. Rev. 37, 159– 170 (2015).

7. Han, B., Jones, C. M., Einstein, E. B., Powell, P. A. & Compton, W. M. Use of Medications for Alcohol Use Disorder in the US: Results From the 2019 National Survey on Drug Use and Health. JAMA Psychiatry 78, 922–924 (2021).

8. Witkiewitz, K., Litten, R. Z. & Leggio, L. Advances in the science and treatment of alcohol use disorder. Sci. Adv. 5, eaax4043 (2019).

9. Leclercq, S. et al. Intestinal permeability, gut-bacterial dysbiosis, and behavioral markers of alcohol-dependence severity. Proc. Natl. Acad. Sci. 111, E4485–E4493 (2014).

10. Addolorato, G. et al. Gut microbiota compositional and functional fingerprint in patients with alcohol use disorder and alcohol-associated liver disease. Liver Int. 40, 878–888 (2020).

11. Day, A. W. & Kumamoto, C. A. Gut Microbiome Dysbiosis in Alcoholism: Consequences for Health and Recovery. Front. Cell. Infect. Microbiol. 12, (2022).

12. Mutlu, E. A. et al. Colonic microbiome is altered in alcoholism. Am. J. Physiol. - Gastrointest. Liver Physiol. 302, G966–G978 (2012).

13. Piacentino, D. et al. Gut microbial diversity and functional characterization in people with alcohol use disorder: A case-control study. PLOS ONE 19, e0302195 (2024).

14. Ames, N. J. et al. Longitudinal gut microbiome changes in alcohol use disorder are influenced by abstinence and drinking quantity. Gut Microbes 11, 1608–1631 (2020).

15. Seo, B. et al. Roseburia spp. Abundance Associates with Alcohol Consumption in Humans and Its Administration Ameliorates Alcoholic Fatty Liver in Mice. Cell Host Microbe 27, 25–40.e6 (2020).

16. Leclercq, S., Schwarz, M., Delzenne, N. M., Stärkel, P. & de Timary, P. Alterations of kynurenine pathway in alcohol use disorder and abstinence: a link with gut microbiota, peripheral inflammation and psychological symptoms. Transl. Psychiatry 11, 503 (2021).

17. Litwinowicz, K., Choroszy, M. & Waszczuk, E. Changes in the composition of the human intestinal microbiome in alcohol use disorder: a systematic review. Am. J. Drug Alcohol Abuse 46, 4–12 (2020).

18. Litwinowicz, K. & Gamian, A. Microbiome Alterations in Alcohol Use Disorder and Alcoholic Liver Disease. Int. J. Mol. Sci. 24, 2461 (2023).

19. Dubinkina, V. B. et al. Links of gut microbiota composition with alcohol dependence syndrome and alcoholic liver disease. Microbiome 5, 141 (2017).

20. Smirnova, E. et al. Fecal Microbiome Distinguishes Alcohol Consumption From Alcoholic Hepatitis But Does Not Discriminate Disease Severity. Hepatology 72, 271 (2020).

21. Carbia, C. et al. The Microbiome-Gut-Brain axis regulates social cognition & craving in young binge drinkers. eBioMedicine 89, (2023).

22. Piacentino, D. et al. Gut microbial diversity and functional characterization in people with alcohol use disorder: A case-control study. PLOS ONE 19, e0302195 (2024).

23. Rao, R. Endotoxemia and gut barrier dysfunction in alcoholic liver disease†. Hepatology 50, 638 (2009).

24. Elamin, E., et al. Ethanol Impairs Intestinal Barrier Function in Humans through Mitogen Activated Protein Kinase Signaling: A Combined In Vivo and In Vitro Approach. PLoS ONE 9, e107421 (2014).

25. Proskynitopoulos, P. J. et al. The effect of alcohol withdrawal therapy on gut microbiota in alcohol use disorder and its link to inflammation and craving. Alcohol Clin. Exp. Res. 49, 1912–1923 (2025).

26. Vatsalya, V. et al. The Beneficial Effects of Lactobacillus GG Therapy on Liver and Drinking Assessments in Patients with Moderate Alcohol-Associated Hepatitis. Am. J. Gastroenterol. 118, 1457–1460 (2023).

27. Bajaj, J. S. et al. A Randomized Clinical Trial of Fecal Microbiota Transplant for Alcohol Use Disorder. Hepatology 73, 1688–1700 (2021).

28. Grodin, E. N., Burnette, E. M., Rodriguez, C., Fulcher, J. A. & Ray, L. A. The gut microbiome in alcohol use disorder and alcohol-associated liver disease: A systematic review of clinical studies. Alcohol Clin. Exp. Res. 48, 1221–1242 (2024).

29. Chen, L., Zhu, Y., Hou, X., Yang, L. & Chu, H. The Role of Gut Bacteria and Fungi in Alcohol-Associated Liver Disease. Front. Med. 9, 840752 (2022).

30. Garrido-Mesa, N., Zarzuelo, A. & Gálvez, J. Minocycline: far beyond an antibiotic. Br. J. Pharmacol. 169, 337–352 (2013).

31. Asadi, A. et al. Minocycline, focus on mechanisms of resistance, antibacterial activity, and clinical effectiveness: Back to the future. J. Glob. Antimicrob. Resist. 22, 161–174 (2020).

32. Leigh, S.-J., Kaakoush, N. O., Westbrook, R. F. & Morris, M. J. Minocycline-induced microbiome alterations predict cafeteria diet-induced spatial recognition memory impairments in rats. Transl. Psychiatry 10, 1–13 (2020).

33. Pechacek, K. M., Reck, A. M., Frankot, M. A. & Vonder Haar, C. Minocycline Fails to Treat Chronic Traumatic Brain Injury-Induced Impulsivity and Attention Deficits. Exp. Neurol. 348, 113924 (2022).

34. Schmidt, E. K. A., Raposo, P. J. F., Torres-Espin, A., Fenrich, K. K. & Fouad, K. Beyond the lesion site: minocycline augments inflammation and anxiety-like behavior following SCI in rats through action on the gut microbiota. J. Neuroinflammation 18, 144 (2021).

35. Wong, M.-L. et al. Inflammasome signaling affects anxiety- and depressive-like behavior and gut microbiome composition. Mol. Psychiatry 21, 797–805 (2016).

36. Hsu, C.-N., et al. Altered Gut Microbiota and Its Metabolites in Hypertension of Developmental Origins: Exploring Differences between Fructose and Antibiotics Exposure. Int. J. Mol. Sci. 22, 2674 (2021).

37. Wang, Y.-L. et al. Microglial activation mediates chronic mild stress-induced depressive- and anxiety-like behavior in adult rats. J. Neuroinflammation 15, 21 (2018).

38. Agrawal, R. G., Hewetson, A., George, C. M., Syapin, P. J. & Bergeson, S. E. Minocycline reduces ethanol drinking. Brain. Behav. Immun. 25, S165–S169 (2011).

39. Crabbe, J. C. et al. Tetracycline derivatives reduce binge alcohol consumption in High Drinking in the Dark mice. Brain Behav. Immun. - Health 4, 100061 (2020).

40. Gajbhiye, S. V., Tripathi, R. K., Salve, B., Petare, A. & Potey, A. V. Evaluation of effect of minocycline on rewarding potential and alcohol relapse in place preference model in mice. Neurosci. Lett. 649, 28–33 (2017).

41. Yang, Q. et al. Chronic minocycline treatment exerts antidepressant effect, inhibits neuroinflammation, and modulates gut microbiota in mice. Psychopharmacology (Berl.) 237, 3201–3213 (2020).

42. Yao, H. et al. Chronic ethanol exposure induced anxiety-like behaviour by altering gut microbiota and GABA system. Addict. Biol. 27, e13203 (2022).

43. Schmidtner, A. K. et al. Minocycline alters behavior, microglia and the gut microbiome in a trait-anxiety-dependent manner. Transl. Psychiatry 9, 1–12 (2019).

44. Thompson, K. G. et al. Minocycline and Its Impact on Microbial Dysbiosis in the Skin and Gastrointestinal Tract of Acne Patients. Ann. Dermatol. 32, 21–30 (2020).

45. Zaura, E. et al. Same Exposure but Two Radically Different Responses to Antibiotics: Resilience of the Salivary Microbiome versus Long-Term Microbial Shifts in Feces. mBio 6, e01693–15 (2015).

46. Husain, M. I. et al. Minocycline as an adjunct for treatment-resistant depressive symptoms: A pilot randomised placebo-controlled trial. J. Psychopharmacol. (Oxf.) 31, 1166– 1175 (2017).

47. Emadi-Kouchak, H. et al. Therapeutic effects of minocycline on mild-to-moderate depression in HIV patients: a double-blind, placebo-controlled, randomized trial. Int. Clin. Psychopharmacol. 31, 20–26 (2016).

48. Hellmann-Regen, J. et al. Effect of Minocycline on Depressive Symptoms in Patients With Treatment-Resistant Depression: A Randomized Clinical Trial. *JAMA Netw*. Open 5, e2230367 (2022).

49. Liu, F. et al. Minocycline supplementation for treatment of negative symptoms in early-phase schizophrenia: A double blind, randomized, controlled trial. Schizophr. Res. 153, 169–176 (2014).

50. Chaudhry, I. B. et al. Minocycline benefits negative symptoms in early schizophrenia: a randomised double-blind placebo-controlled clinical trial in patients on standard treatment. J. Psychopharmacol. (Oxf.) 26, 1185–1193 (2012).

51. Deakin, B. et al. The benefit of minocycline on negative symptoms of schizophrenia in patients with recent-onset psychosis (BeneMin): a randomised, double-blind, placebo-controlled trial. Lancet Psychiatry 5, 885–894 (2018).

52. Arout, C. A., Waters, A. J., MacLean, R. R., Compton, P. & Sofuoglu, M. Minocycline does not affect experimental pain or addiction-related outcomes in opioid maintained patients. Psychopharmacology (Berl.) 236, 2857–2866 (2019).

53. Huang, H., Zou, Z. & Chen, B. Efficacy of Minocycline in Depression: A Systematic Review and Meta-analysis. Clin. Neuropharmacol. 48, 1–6 (2025).

54. American Psychiatric Association. Diagnostic and Statistical Manual of Mental Disorders (5th Ed.). (American Psychiatric Association, Arlington, VA, 2013).

55. Segal-Isaacson, C. J., Wylie-Rosett, J. & Gans, K. M. Validation of a Short Dietary Assessment Questionnaire: The Rapid Eating and Activity Assessment for Participants Short Version (REAP-S). Diabetes Educ. 30, 774–781 (2004).

56. Cerwinske, L. A., Rasmussen, H. E., Lipson, S., Volgman, A. S. & Tangney, C. C. Evaluation of a dietary screener: the M editerranean E ating P attern for A mericans tool. J. Hum. Nutr. Diet. 30, 596–603 (2017).

57. Sobell, L. C. & Sobell, M. B. Timeline follow-back: A technique for assessing self-reported alcohol consumption. in Measuring Alcohol Consumption: Psychosocial and Biological Methods. (Humana Press, Totowa, NJ, 1992).

58. First, M. B. & Williams, J. B. W. Structured Clinical Interview for DSM-5 Disorders -Reserch Version (SCID-5-RV). (American Psychiatric Association, Arlington, VA, 2015).

59. Bohn, M. J., Krahn, D. D. & Staehler, B. A. Development and initial validation of a measure of drinking urges in abstinent alcoholics. Alcohol. Clin. Exp. Res. 19, 600–606 (1995).

60. Holm, J. B. et al. Ultrahigh-Throughput Multiplexing and Sequencing of >500-Base-Pair Amplicon Regions on the Illumina HiSeq 2500 Platform. mSystems 4, e00029–19 (2019).

61. Quast, C. et al. The SILVA ribosomal RNA gene database project: improved data processing and web-based tools. Nucleic Acids Res. 41, D590–D596 (2012).

62. Wang, Q., Garrity, G. M., Tiedje, J. M. & Cole, J. R. Naïve Bayesian Classifier for Rapid Assignment of rRNA Sequences into the New Bacterial Taxonomy. Appl. Environ. Microbiol. 73, 5261–5267 (2007).

63. McMurdie, P. J. & Holmes, S. phyloseq: An R Package for Reproducible Interactive Analysis and Graphics of Microbiome Census Data. PLOS ONE 8, e61217 (2013).

64. Bates, D., Mächler, M., Bolker, B. & Walker, S. Fitting Linear Mixed-Effects Models Using **lme4**. J. Stat. Softw. 67, (2015).

65. Lenth, R. V., et al. emmeans: Estimated Marginal Means, aka Least-Squares Means. (2026).

66. Gloor, G. B., Macklaim, J. M., Pawlowsky-Glahn, V. & Egozcue, J. J. Microbiome Datasets Are Compositional: And This Is Not Optional. Front. Microbiol. 8, 2224 (2017).

67. Nickols, W. A. et al. MaAsLin 3: Refining and extending generalized multivariable linear models for meta-omic association discovery. Preprint at 10.1101/2024.12.13.628459 (2024).

68. Lin, S.-K. K. et al. Dysbiosis and depression: A study of gut microbiota alterations and functional pathways in antidepressant-naïve mood disorder patients. Transl. Psychiatry 15, 290 (2025).

69. Louis, P. & Flint, H. J. Formation of propionate and butyrate by the human colonic microbiota. Environ. Microbiol. 19, 29–41 (2017).

70. Zhu, F. et al. Metagenome-wide association of gut microbiome features for schizophrenia. Nat. Commun. 11, 1612 (2020).

71. Smirnova, E. et al. FECAL MICROBIOME DISTINGUISHES ALCOHOL CONSUMPTION FROM ALCOHOLIC HEPATITIS BUT DOES NOT DISCRIMINATE DISEASE SEVERITY. Hepatol. Baltim. Md 72, 271–286 (2020).

72. Davis, J. A. et al. The associations of butyrate-producing bacteria of the gut microbiome with diet quality and muscle health. Gut Microbiome 2, e2 (2021).

73. Gurwara, S. et al. Alcohol use alters the colonic mucosa-associated gut microbiota in humans. Nutr. Res. N. Y. N 83, 119–128 (2020).

74. Kuo, C.-H., El-Omar, E., Kao, C.-Y., Lin, J.-T. & Wu, C.-Y. Compositional and Metabolomic Shifts of the Gut Microbiome in Alcohol-Related Liver Disease. J. Gastroenterol. Hepatol. 40, 2176–2189 (2025).

75. Zhong, X. et al. Streptococcus, the Predominant Bacterium to Predict the Severity of Liver Injury in Alcoholic Liver Disease. Front. Cell. Infect. Microbiol. 11, 649060 (2021).

76. Misera, A. et al. Comparative analysis of gut microbiota in major depressive disorder and schizophrenia during hospitalisation - the case-control, post hoc study. Psychoneuroendocrinology 171, 107208 (2025).

77. Chen, J. et al. An expansion of rare lineage intestinal microbes characterizes rheumatoid arthritis. Genome Med. 8, 43 (2016).

78. Thirion, F. et al. Alteration of Gut Microbiome in Patients With Schizophrenia Indicates Links Between Bacterial Tyrosine Biosynthesis and Cognitive Dysfunction. Biol. Psychiatry Glob. Open Sci. 3, 283–291 (2023).

79. O’Hare, M. A. et al. The saNeuroGut Initiative: Investigating the Gut Microbiome and Symptoms of Anxiety, Depression, and Posttraumatic Stress. Neuroimmunomodulation 32, 1–15 (2025).

80. Gomez-Arango, L. F. et al. Low dietary fiber intake increases Collinsella abundance in the gut microbiota of overweight and obese pregnant women. Gut Microbes 9, 189–201 (2018).

81. Sadoughi, B., Schneider, D., Daniel, R., Schülke, O. & Ostner, J. Aging gut microbiota of wild macaques are equally diverse, less stable, but progressively personalized. Microbiome 10, 95 (2022).

82. Lopetuso, L. R., Scaldaferri, F., Petito, V. & Gasbarrini, A. Commensal Clostridia: leading players in the maintenance of gut homeostasis. Gut Pathog. 5, 23 (2013).

83. Vacca, M. et al. The Controversial Role of Human Gut Lachnospiraceae. Microorganisms 8, 573 (2020).

84. Kim, Y.-J., Jung, D.-H. & Park, C.-S. Important roles of Ruminococcaceae in the human intestine for resistant starch utilization. Food Sci. Biotechnol. 33, 2009–2019 (2024).

85. Coello, K. et al. Affective disorders impact prevalence of Flavonifractor and abundance of Christensenellaceae in gut microbiota. Prog. Neuropsychopharmacol. Biol. Psychiatry 110, 110300 (2021).

86. Waters, J. L. & Ley, R. E. The human gut bacteria Christensenellaceae are widespread, heritable, and associated with health. BMC Biol. 17, 83 (2019).

87. Jaroszewski, J., Mamun, N. & Czaja, K. Bidirectional Interaction between Tetracyclines and Gut Microbiome. Antibiotics 12, 1438 (2023).

88. Singh, S. B., Carroll-Portillo, A. & Lin, H. C. Desulfovibrio in the Gut: The Enemy within? Microorganisms 11, 1772 (2023).

89. Yang, G. et al. Discovery of an ene-reductase for initiating flavone and flavonol catabolism in gut bacteria. Nat. Commun. 12, 790 (2021).

90. Ju, T., Kong, J. Y., Stothard, P. & Willing, B. P. Defining the role of Parasutterella, a previously uncharacterized member of the core gut microbiota. ISME J. 13, 1520–1534 (2019).

91. Moura, I. B. et al. Profiling the Effects of Systemic Antibiotics for Acne, Including the Narrow-Spectrum Antibiotic Sarecycline, on the Human Gut Microbiota. Front. Microbiol. 13, 901911 (2022).

92. Barb, J. J. et al. Dietary intake, quality, and assessment tools in individuals with problematic alcohol use: a scoping review and meta-analysis. Transl. Psychiatry 16, 51 (2026).

